# A single particle analysis method for detecting membrane remodelling and curvature sensing

**DOI:** 10.1101/2024.04.09.588706

**Authors:** Adeline Colussi, Leonardo Almeida-Souza, Harvey T McMahon

**Author notes:** Correspondence should be address to LA-S and H.T.M. These authors have jointly supervised this work (Leonardo Almeida-Souza, Harvey T McMahon).

## Abstract

In biology, shape and function are related. Therefore, it is important to understand how membrane shape is generated, stabilised and sensed by proteins and how this relates to organelle function. Here we present an assay that can detect curvature preference and membrane remodelling using free-floating liposomes using protein concentrations in a physiologically relevant ranges. The assay reproduced known curvature preferences of BAR domains and allowed the discovery of high curvature preference for the PH domain of AKT and the FYVE domain of HRS. In addition, our method reproduced the membrane vesiculation activity of the ENTH domain of Epsin1 and showed similar activity for the ANTH domains of PiCALM and Hip1R. Finally, we found that the curvature sensitivity of the N-BAR domain of Endophilin inversely correlates to membrane charge and that deletion of its N-terminal amphipathic helix increased its curvature specificity. Thus, our method is a generally applicable qualitative method for assessing membrane curvature sensing and remodelling by proteins.

## Introduction

Eukaryotic cells are characterised by membranes with varied and dynamic compositions, topologies and morphologies, ranging from elongated tubules and highly curved vesicles to flat membrane areas. Local modulation of the curvature and composition of membranes generate platforms able to recruit and activate proteins that orchestrate, in space and time, multiple cellular activities. If membrane curvature is to have a function, then proteins must be able to sense it. Moreover, there are many proteins in the cell that help create highly curved vesicular trafficking intermediates and are involved in both fission and fusion of these structures. Therefore, assays capable of measuring curvature preferences and membrane remodelling will be valuable tools to understand protein function at membranes.

To accurately determine curvature preferences of proteins, one would need to generate liposomes of homogeneous size and test for protein binding. Unfortunately, the main method for making liposomes of defined sizes is by passing large liposomes through filters of defined pore sizes (MacDonald et al., 1991) which results in liposomes with an upper limit of size, but a range of liposomes that are smaller than the diameter of the pores (Kunding et al., 2008). To overcome this limitation, one can find alternative methods to generate specific curvatures or develop techniques that determine which liposome sizes specific proteins bind to in a heterogeneous liposome population. Using these principles, a few methods have been developed to study curvature sensitivity (Hatzakis et al., 2009; Lu et al., 2023; Jin et al., 2022; Hsieh et al., 2012; Sorre et al., 2012) (see Discussion).

Here we present a single particle detection method where the hydrodynamic radii of liposomes are calculated from their Brownian motion using the Stokes-Einstein equation. Our method is implemented on a NanoSight instrument controlled by the Nanoparticle Tracking Analysis (NTA) software (commercialised by Malvern). Our method allows the qualitative determination of curvature preference of proteins by identifying the subpopulation of liposomes to which a protein is bound and to study membrane remodelling properties of proteins. Using this technique, we can reproduce curvature sensitivities of different Bin/Amphiphysin/Rvs (BAR) domain-containing proteins and the membrane remodelling activity of the Epsin ENTH (Epsin N-terminal Homology) domain. By applying this method, we identify new curvature-sensitive lipid-binding domains and domains with remodelling activity. Moreover, this technique allows us to better understand the mechanism of curvature sensitivity and generation by the N-BAR (N-terminal helix and Bin/Amphiphysin/Rvs) domain of Endophilin.

## Results

### A method to detect liposome subpopulations

Our method tracks single liposomes, infers their size from their diffusion coefficient, identifies fluorescent protein-bound liposomes as fluorescent objects, and compares the particle-size statistics of protein-bound and unbound liposomes to assess curvature-dependent protein binding. For that, we have used the NanoSight instrument from Malvern to record movies of liposomes in order to analyse the Brownian motion of individual particles using the supplied NTA software, which implements the Stokes-Einstein equation.

Liposomes are made to flow through a specially designed glass chamber with associated laser source, mounted on an upright microscope (Fig. 1a,b). A laser beam illuminates the sample perpendicular to the objective, enabling the trajectory of liposomes freely moving in solution to be tracked using diffracted light. Short movies (120s at 25fps) are collected and many individual particles are tracked using the proprietary NTA software, where the hydrodynamic radius for each particle is calculated. The data are binned in five nanometer intervals and results plotted as a size distribution plot (Fig. 1b). By adding a long-pass filter to the optical path, only particles containing fluorophores are sized (Fig. 1a). More detailed background on the NanoSight instrument and how the methods works can be found in Supplementary Note 1.

**Figure 1.**
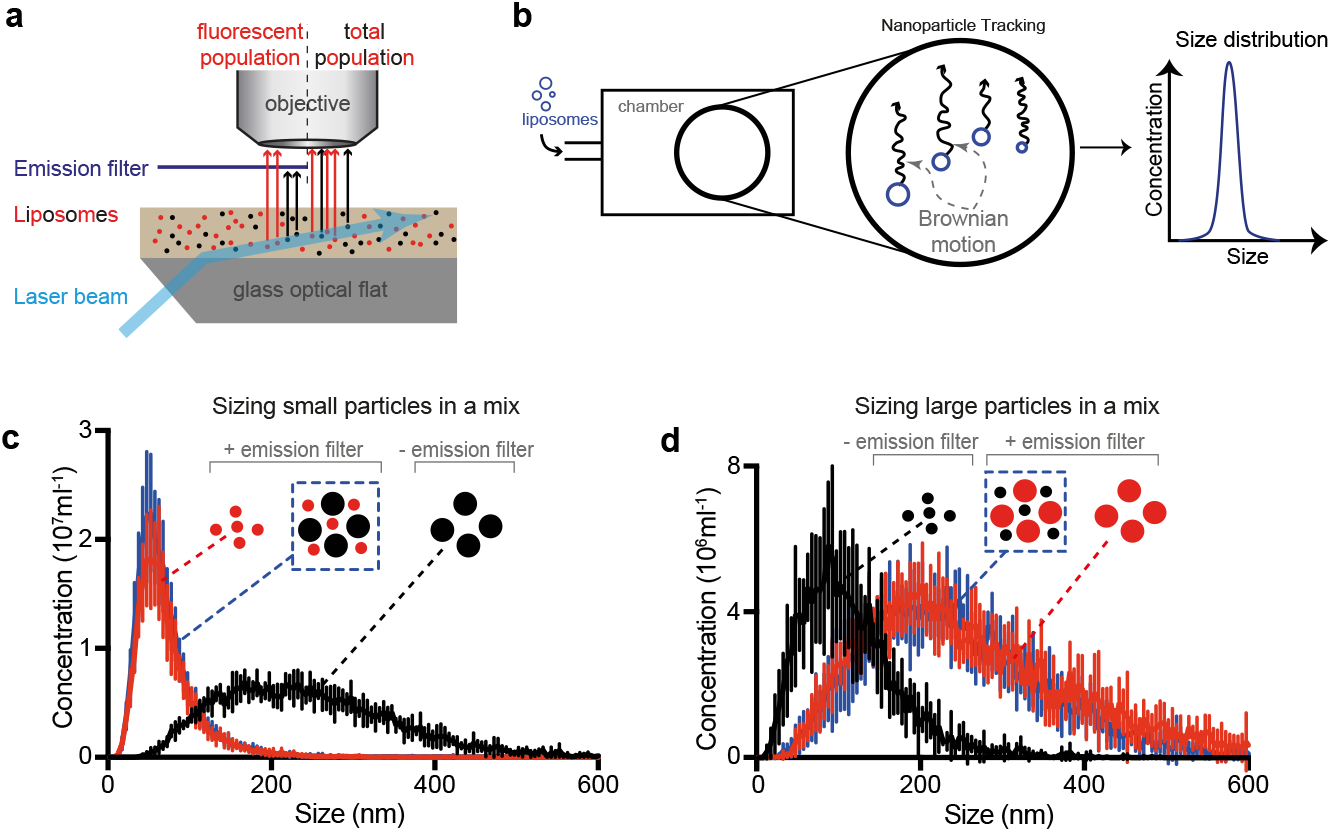
Working principle and validation. **a**. Scheme of how liposomes are detected using diffraction or fluorescence modes. Upon exiting the glass optical flat, the laser beam is refracted through the liquid sample and diffracted by particles in the sample. Introduction of an emission filter that blocks diffracted light allows detection of fluorescent particles only. **b**. Scheme of setup. Liposome samples (blue) are introduced in the sample chamber where they freely move due to Brownian motion. Each particle is tracked and its size is calculated according to Stokes-Einstein equation. Size distribution is given as the concentration of particles in each 5 nm bin. **c-d**. Sizing of a fluorescent subpopulation of liposomes is unaffected by the presence of non-fluorescent liposomes. Small (c) or large (d) fluorescent liposomes were sized in the absence (red) or presence (blue) of a population of non-fluorescent liposomes of a different size (black). Black trace shows the size distribution of the non-fluorescent liposomes population detected by diffraction (no emission filter).

To illustrate the sizing capabilities of our setup, we used calibration beads of defined sizes (100nm and 216nm) and compared the size determination accuracy of our setup with Dynamic Light Scattering (DLS) - an established particle sizing method. Similar to what has been described before (Filipe et al., 2010), The particle tracking analysis was both more precise and accurate than DLS, as it could measure bead sizes more exactly and with a sharper distribution (Supplementary Fig. 1a,c). Importantly, NTA could clearly distinguish two populations of beads when mixed together (Supplementary Fig. 1b,c), something we could not achieve with DLS using the same sample.

We then tested the capacity of our setup to measure liposome sizes and to distinguish subpopulations of fluorescent liposomes mixed with a non-fluorescent liposome population (Fig. 1c,d). We were able to size a subpopulation of small or large fluorescent liposomes alone (Fig. 1c,d - red traces) or mixed together with non-fluorescent liposomes of a different size (Fig. 1c,d - blue traces).

When measuring the same population of liposomes by diffraction and fluorescence, we noticed that the smaller particles were detected much more efficiently by fluorescence (Supplementary Fig. 1d). This is due to the fact that liposomes are weakly diffracting particles and therefore, smaller liposomes are harder to detect by diffraction alone. This hidden fraction was especially obvious with liposomes extruded through 50nm pores. Therefore, we implemented a correction factor to account for this discrepancy when measuring diffraction of liposomes extruded using 50nm filters. (Supplementary Fig. 1e,f).

### A method to measure curvature preference

To accomplish curvature preference measurements, the principle of our method is as follows: Using a population of liposomes of various sizes mixed with a fluorescently labelled protein (or protein domain) of interest, we were be able to determine the curvature preference of said protein by directly comparing size distributions of the total population of liposomes versus fluorescent protein-associated populations (Fig. 2a). To facilitate the visualisation of the results, curvature preferences are displayed as box plots where the middle line represents the peak of the distributions (mode) and the upper and bottom box boundaries represent 50% of the data above and below the mode, respectively (Fig. 2b). The ability to measure fluorescence sets this method apart from other bulk methods as one can detect the sub-population of protein-bound liposomes and compare it to the total population of liposomes. Thus, while co-sedimentation assays with liposomes filtered to different sizes give a view of the tendency of protein to prefer either large or small liposomes(Peter et al., 2004; Henne et al., 2007), our method gives a detailed view of preferences within a filtered population.

**Figure 2.**
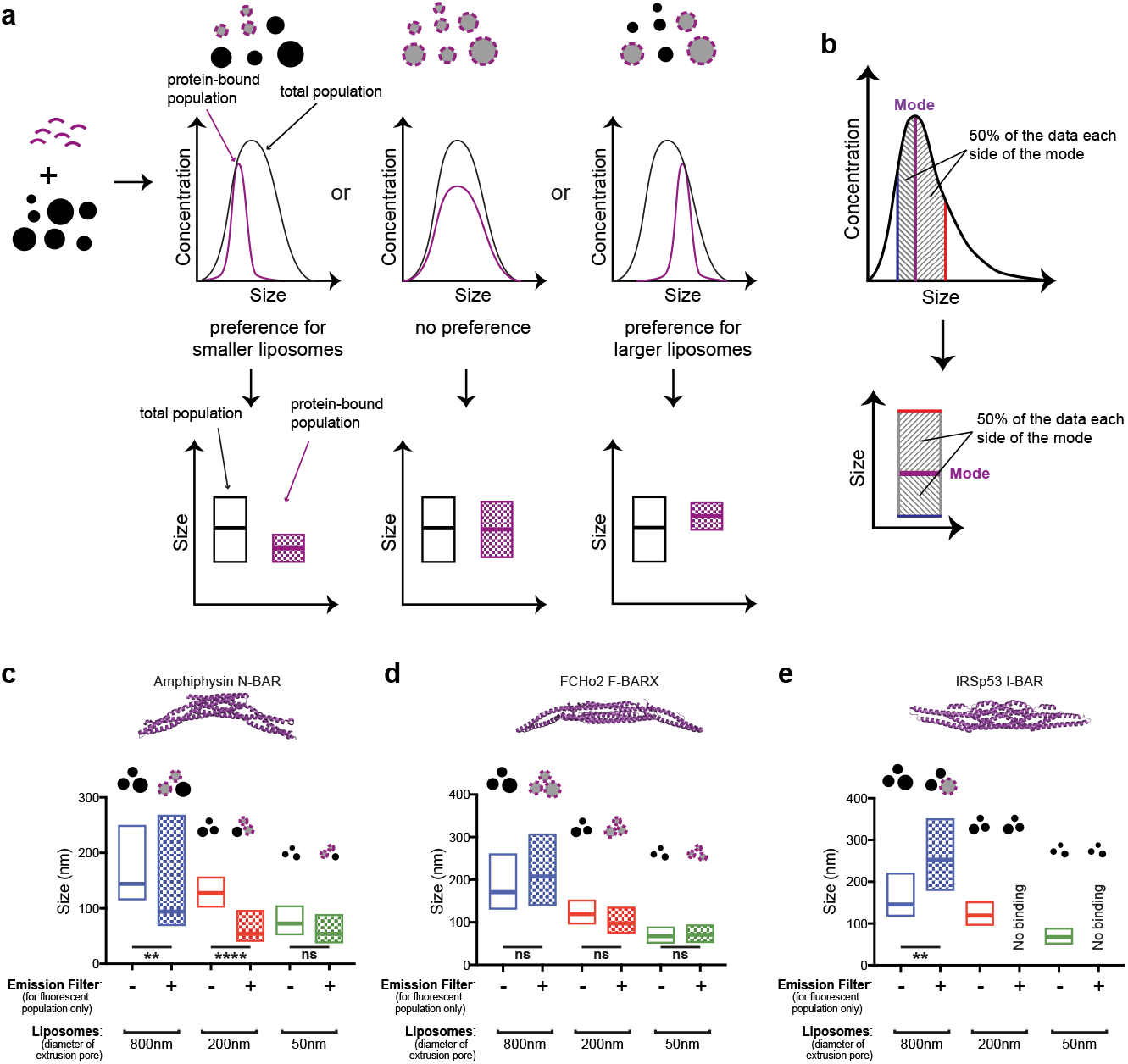
Method principle and validation using BAR domains. **a**. Scheme for the curvature sensitivity assay. Fluorescent protein (purple) is incubated with non-fluorescent liposomes (black circles) and measured in our SETUP. Comparison between the size distributions of all liposomes (black curve) with the fluorescent signal of the protein-bound sub-population (purple) indicates whether there is a curvature preference for small liposomes, large liposomes or an absence of curvature preference. The data can then be displayed using boxplots as shown in (b), the pattern-filled box represents the fluorescent, protein-bound subpopulation. **b**. Box plots display of results. Box plots show the mode of the size distribution as the middle line while boundaries represent 50% of the data on each side of the mode. **c-e**. Measurents for (c) Amphiphysin N-BAR (pdb: 1uru), (d) FCHo2 F-BAR (pdb: 2v0o) and (e) IRSP53 I-BAR (pdb: 1y2o). **c**. Amphiphysin N-BAR shows specificity for high curvatures. **d**. FCHo2 F-BAR is insensitive to curvature and binds all sizes of liposomes equally. **e**. IRSp53 I-BAR preferentially binds to the largest liposomes and does not bind to smaller liposomes extruded at 200 or 50 nm.Boxplots depict mode values (middle line) -/+ 50% data on each side of the mode (indicated by bottom and top lines). n=3, ns = non-significant, p < 0.01 (^**^), p < 0.0001 (^****^). p-values tested using ANOVA followed by Bonferroni correction for multiple comparisons.

The structures of several BAR domains have been determined and are seen to have different intrinsic curvatures which translate into different membrane curvature preferences (Peter et al., 2004; Millard et al., 2005; Prévost et al., 2015; Barooji et al., 2016; Henne et al., 2007; Simunovic et al., 2019), and thus these can act as good controls for validating our method. For this initial validation we used three types of BAR domain: Amphiphysin N-BAR domain, which has been described to bind to small liposomes (Peter et al., 2004); FCHo2 (F-BAR domain only protein 2) F-BAR (Fes/Cip4 homology Bin/Amphiphysin/ Rvs) domain, which shows no curvature preference (Henne et al., 2007); and IRSp53 (Insulin receptor substrate protein of 53 kDa) I-BAR (inverted Bin/Amphiphysin/Rvs) domain which binds to larger (*i*.*e*. flatter, or negatively curved) membranes (Millard et al., 2005; Prévost et al., 2015; Barooji et al., 2016). Monomeric, enhanced, superfolder green fluorescent protein (mesfGFP (Zhang et al., 1996; Zacharias et al., 2002; von Stetten et al., 2012a)) tagged BAR domains were added to three different liposome populations extruded through 800nm, 200nm and 50nm filters and the sizes of the total liposome population and the protein bound subpopulation were compared. As shown in figures 2c-e, the curvature preference for all BAR domains was correctly determined by our method (see also Supplementary Fig. 2a-c). Amphiphysin N-BAR preferentially bound to the smallest available liposomes (Fig. 2c, Supplementary Fig. 2a), while FCHo2 F-BAR was insensitive to membrane curvature (Fig. 2d, Supplementary Fig. 2b). IRSp53 I-BAR was found exclusively on the largest liposomes of the 800nm-extruded sample and did not bind liposomes extruded with smaller pore sizes (Fig. 2e, Supplementary Fig. 2c). By electron microscopy, BAR domain proteins have been seen to tubulate membranes at micromolar concentrations(Mattila et al., 2007; Ambroso et al., 2014; Henne et al., 2007). As a single particle method, our assay requires nanomolar concentrations of fluorescently-labelled proteins (∼2nM) and as shown in Supplementary Figure 2d, under these concentrations, Amphiphysin did not affect the size distribution of the total liposome population, confirming that the smaller liposomes fluorescently labelled by Amphiphysin came from curvature sensing rather than membrane remodelling. Under those conditions we therefore could detect curvature sensitivity without causing morphological alterations to the population of liposomes.

Having established our method can measure the curvature sensitivity of proteins, we next applied it to a set of proteins from different families of lipid binding domains. In an initial screen using Folch lipids, many domains did not bind, therefore we tuned the lipid compositions for each domain according to information available in the literature (Supplementary Table 1). Using optimal conditions for binding, we found that the pleckstrin homology (PH) domain of AKT (protein kinase B) and the FYVE (Fab1, YOTB, Vac1, EEA1) domain of HRS (hepatocyte growth factor-regulated tyrosine kinase substrate) are curvature sensitive (Fig. 3a-b, Supplementary Fig. 3a-b), binding preferentially to smaller liposomes. To our knowledge, this is the first time a specific curvature preference has been described for these particular domains. On the other hand, the C1 domain of PKCß2 (protein kinase C beta) and the FERM (4.1, ezrin/radixin/moesin) domain of Talin1 did not show any curvature specificity and could bind all liposome sizes (Fig. 3c-d, Supplementary Fig. 3c-d). The results for the lipid binding domains tested and the corresponding lipid compositions are summarised in Supplementary Table 1. Taken together, our results showed that our method is capable of determining curvature preferences for a multitude of structurally diverse lipid binding domains, once binding conditions have been established.

**Figure 3.**
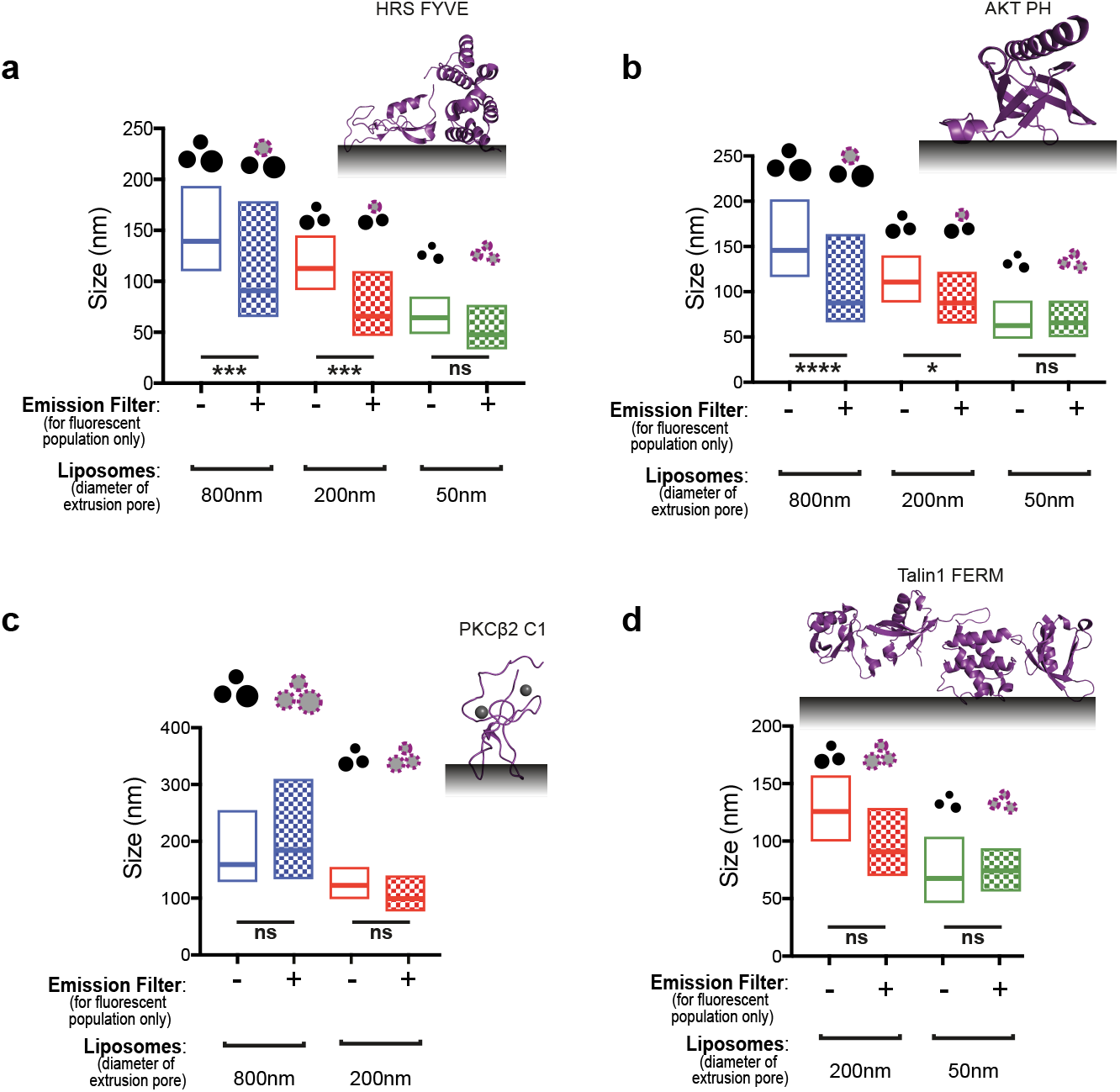
Curvature preference for diverse lipid-binding domains. **a**.FYVE domain of HRS (pdb: 1dvp) displays preference for higher curvatures (smaller liposomes). HRS FYVE domain preferentially binds to the subpopulation of small liposomes on the 800 nm or 200 nm extruded samples and all sizes on the smaller, 50nm extruded sample. **b**. PH domain of AKT (pdb: 1unq) displays preference for higher curvatures (smaller liposomes). AKT PH domain preferentially binds to the subpopulation of small liposomes on the 800 nm or 200 nm extruded samples and all sizes on the smaller, 50 nm extruded sample. **c**. C1B domain of PKCβ2 (pdb: 3pfq) displays no curvature specificity and binds all sizes of liposomes in 800 and 200 nm extruded samples. **d**. FERM domain of Talin1 (pdb: 3ivf) displays no curvature specificity and binds all sizes of liposomes indifferently in all samples. Boxplots depict mode values (middle line) -/+ 50% data on each side of the mode (indicated by bottom and top lines). n=3, ns = non-significant, p < 0.05 (^*^), p<0.001 (^***^), p<0.0001 (^****^). p-values tested using ANOVA followed by Bonferroni correction for multiple comparisons.

### A method to detect membrane remodelling

The capacity of our setup to accurately size particles in solution opens the possibility that this method can also monitor membrane-remodelling events. To test this, we took advantage of the liposome remodelling properties of the ENTH domain of Epsin1. We have previously shown that the Epsin1 ENTH domain can generate smaller vesicles (vesiculation) by inserting its N-terminal amphipathic helix into the outer membrane leaflet (Ford et al., 2002a). Given the sizing capabilities of our setup, we reasoned that we should be able to detect vesiculation by measuring a decrease of average liposome size as well as an increase of particle concentration (Fig. 4a).

**Figure 4.**
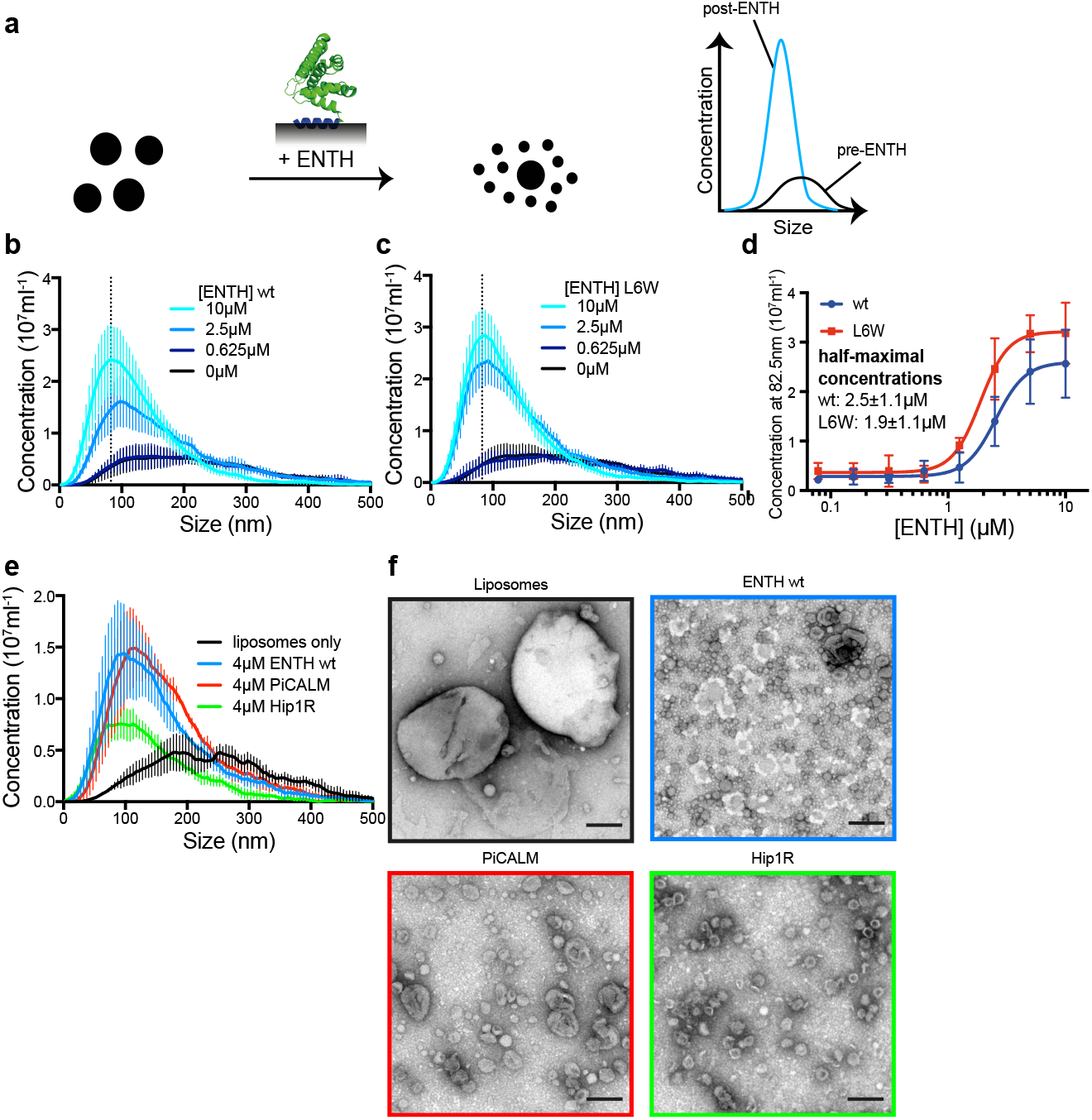
Monitoring protein-induced vesiculation. **a**. Scheme for the assay to detect membrane vesiculation. Incubation of liposomes with ENTH (Epsin N-terminal homology domain, pdb: 1h0a) induces formation of smaller liposomes by vesiculation. This is observed by a reduction in mean liposome size and an increase in concentration of particles. **b-c**. Vesiculation by ENTH wild-type (wt, b) or the hyperactive L6W mutant (c) after incubation of liposomes with different ENTH concentrations. **d**. Corresponding dose-response curves for vesiculation by ENTH wt or L6W. The concentration of particles in the bin centred around 82.5 nm (marked as a dotted line in (b,c)) was used as a marker of vesiculation efficiency. **e**. Comparison of vesiculation efficiency between ENTH, PiCALM ANTH and Hip1R ANTH domains. PiCALM and Hip1R ANTH both vesiculate with PiCALM producing larger vesicles than ENTH and Hip1R fewer vesicles. **f**. Negative-stain EM of liposomes before (black frame) and after incubation with 4 μM ENTH (blue frame), PiCALM ANTH (red frame) or Hip1R ANTH (green frame). Scale bar = 200 nm.

To test this possibility, fluorescent liposomes were sized in our setup before and after incubation with increasing concentrations of unlabelled Epsin1 ENTH domain. For comparison, we also sized liposomes incubated with the Epsin1 ENTH domain mutant L6W, that previously was shown to exhibit enhanced vesiculating activity (Ford et al., 2002b). In accordance with its vesiculation properties, liposomes incubated with Epsin1 ENTH domain showed a dose dependent decrease in mean size and an increase in particle concentration (Fig. 4b,d). Importantly, the L6W mutant was more efficient and generated more vesicles than wild-type ENTH (Fig. 4c,d, p-value = 0.032).

The ANTH (AP180 N-terminal homology) domains of PiCALM (Phosphatidylinositol-binding clathrin assembly protein) and Hip1R (Huntingtin-interacting protein 1-related protein) share structural homology with the ENTH domain. Moreover, the ANTH domain of PiCALM has been shown to induce tubulation (Miller et al., 2015). We therefore tested if ANTH domains could also vesiculate liposomes and observed that both PiCALM and Hip1R ANTH domains did so in a dose dependent manner (Supplementary Fig. 4a-b). Comparison between ENTH, PiCALM and Hip1R ANTH showed that the vesicles generated by PiCALM were slightly larger than ENTH and that Hip1R was vesiculating much less effectively (Fig. 4e). We confirmed these results by electron microscopy (EM) observations of the vesicles generated by these proteins (Fig. 4f).

Thus, our results show that our method is also capable of detecting membrane remodelling events by ENTH domain and reveal, for the first time, that the homologous ANTH domain can also vesiculate membranes.

### A method to differentiate curvature sensing and generation

The concepts of membrane curvature sensing and curvature generation are on a spectrum and are intrinsically linked and hard to distinguish using conventional methods. This is partly because curvature sensors can, by mass action of high protein concentrations, also generate curvature (Stachowiak et al., 2012). As our method allows the clear identification of curvature preference using nanomolar protein concentrations, we applied it to better understand the binding of Endophilin N-BAR to membranes.

First, we studied the influence of membrane binding affinity on curvature preference. For that, we measured the curvature preference of Endophilin N-BAR domain on liposomes of increasing charge density, given that membrane binding is thought to be mediated by positively charged protein residues and anionic phospholipid headgroups. Although Endophilin N-BAR displayed a strong preference for higher curvatures on liposomes with a lower charge density (40% Folch extract in phosphatidylcholine), the curvature sensitivity decreased with increasing liposome charge and disappeared with 80% Folch liposomes (Fig. 5a, Supplementary Fig. 5a).

**Figure 5.**
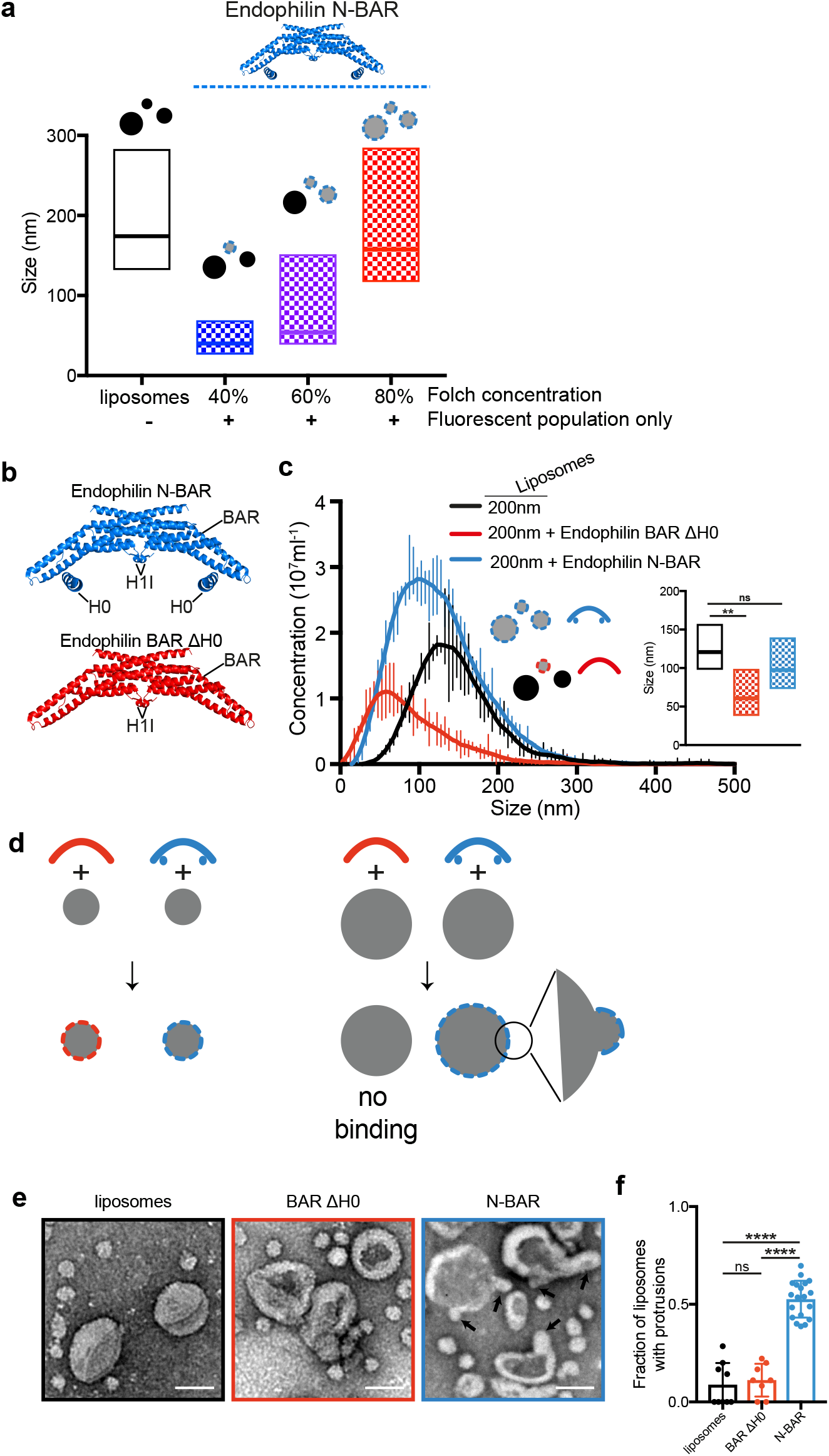
Mechanism of curvature sensing by Endophilin. **a**. Influence of charge on membrane curvature sensing. Increasing the concentration of Folch lipids containing charged phospholipid headgroups in a phosphatidylcholine background reduces the stringency of curvature sensing by Endophilin N-BAR. **b**. Structure of N-BAR and BAR ΔH0 constructs of Endophilin used (pdb: 2c08). The H0 was added manually for illustration purposes as the H0 is not visible in the available Endophilin crystal structures. **c**. Curvature sensitivity of Endophilin N-BAR and BAR ΔH0 to highly charged liposomes (1:1 FolchS/FolchA). Endophilin N-BAR (blue) binds all sizes of 200 nm extruded liposomes (black), whereas Endophilin BAR ΔH0 (red) specifically binds a subpopulation of smaller liposomes. **d**. Model of Endophilin N-BAR and BAR ΔH0 binding to small and large liposomes. Endophilin BAR ΔH0 is specific for small liposomes and does not bind to large liposomes. Endophilin N-BAR binds both small and large liposomes, in that case by inducing high local curvature to which it can bind through the BAR domain. **e**. Negative-stain EM of liposomes before (black frame) and after addition of Endophilin BAR ΔH0 (red frame) or Endophilin N-BAR (blue frame). Endophilin N-BAR induces formation of areas with high local curvature on large liposomes indicated by arrows. Scale bar = 100 nm. **f**. Quantification of the fraction of liposomes showing protrusions in each micrograph for liposomes alone (n=9) or in the presence of Endophilin BAR ΔH0 (n=8) or Endophilin N-BAR (n=19). p-values tested using ANOVA followed by Bonferroni correction for multiple comparisons, ns = non-significant, p<0.0001 (^****^).

Two modes of curvature sensing have been described. Curved protein domains forming large electrostatic contact surfaces, like BAR domains, which can preferentially bind similarly curved membranes (Gallop et al., 2006; Barooji et al., 2016); and amphipathic helices (or other features) inserting into hydrophobic defects which preferentially bind highly curved membranes (Bigay et al., 2003; Drin et al., 2007). Endophilin N-BAR contains both a curved BAR domain and an amphipathic helix H0 (Fig. 5b) and their relative contribution towards curvature sensing is debated (Gallop et al., 2006; Bhatia et al., 2009). Using our method, we compared the curvature sensitivity of Endophilin N-BAR and Endophilin BAR ΔH0 (a construct where the N-terminal amphipathic helix H0 was deleted) on Folch liposomes (Fig. 5c). Whereas Endophilin N-BAR bound liposomes of all sizes with only a slight preference for smaller liposomes (Fig. 5c, blue), deletion of H0 increased curvature sensitivity (Fig. 5c, red). This is not due to vesiculation as neither the size of the whole liposome population after addition of protein (Supplementary Fig. 5b) nor the size of the fluorescent population after longer incubation times changed (Supplementary Fig. 5c).

This increased curvature preference for Endophilin ΔH0 over N-BAR leads to a model (Fig. 5d) by which the BAR domain of Endophilin acts as the curvature sensor and H0 helices induce local deformation of lipid membranes, allowing the BAR domain to bind to liposomes larger than its own curvature. In accordance with this model, liposomes incubated with Endophilin N-BAR at the same lipid:protein ratios used for the curvature preference measurements showed small bulges consistent with local membrane deformations. Importantly, these bulges were absent on liposomes alone or liposomes incubated with Endophilin BAR ΔH0 (Fig. 5e-f).

## Discussion

The method we present here is a simple and powerful tool to study membrane curvature preference of proteins and membrane remodelling. Classically, membrane curvature preferences for proteins have been measured using liposome co-sedimentation or flotation assays (Bigay et al., 2005; Peter et al., 2004). The difficulty to produce homogeneously sized liposomes render these classical techniques suitable only for detecting proteins with more extreme curvature preferences (Kunding et al., 2008; Bhatia et al., 2009). Moreover, the high protein concentrations required for these assays may be a confounding factor as many curvature sensors can deform membranes at high concentrations (Roux et al., 2010; Peter et al., 2004; Mattila et al., 2007; Gallop et al., 2006). The capacity of our method to accurately size thousands of individual liposomes and to detect binding to membranes at nanomolar concentrations of protein represents a significant advance from these bulk assays.

Over the years, methods using solid supported membranes, tethered liposomes and lipid nanotubes have been developed to study membrane curvature (Hatzakis et al., 2009; Hsieh et al., 2012; Jin et al., 2022; Lu et al., 2023; Sorre et al., 2012). The peculiarities of each of these methods make them appropriate for specific applications. For example, lipid nanotubes are remarkable tools to study the effect of protein binding under membrane tension (Simunovic et al., 2017). On the other hand, lipid supported membranes (Zhao et al., 2017) and liposomes attached to a surface are especially suited for visualisation of dynamic processes or the detection of negative membrane curvatures (Hatzakis et al., 2009; Lu et al., 2023). The assay we describe here is a general method that is applicable to proteins with multiple positive curvature preferences using unperturbed, free-floating liposomes. A crucial advantage of our method is that it uses an unmodified, commercially available instrument with an easy-to-use software. This allows groups with access to this instrument to easily test the suitability of our method to answer their research question. On the other hand, the difficulty to modify the instrument or modify the software controlling it may prove a barrier for more complex applications.

Curvature sensing and generation are tightly linked processes and increasing evidence shows that they are part of the same continuum. Several BAR-domains have been shown to be able to both sense and generate tubules or vesicles depending on the concentrations used (Endophilin (Gallop et al., 2006), FCHo2 (Peter et al., 2004), IRSp53 (Mattila et al., 2007; Prévost et al., 2015)). Similarly, Dynamin, the mediator of vesicle scission in multiple endocytic pathways, has been shown to be a curvature sensor at low concentrations (Merrifield et al., 2002; Roux et al., 2010) and a curvature generator at high concentrations (Pucadyil and Schmid, 2009). A study on Amphiphysin N-BAR domain on tubules pulled from a giant unilamellar vesicle (GUV) showed that depending on protein density, Amphiphysin senses curvature and preferentially binds to the narrow tubule (at low protein density) or tubulates the GUV at higher concentrations (Sorre et al., 2012). With different phenomena happening depending on protein concentration, membrane rigidity and tension, obtaining quantitative data on curvature preference has proven difficult until now. Moreover, likely for technical reasons, there is a bias in the field towards the study of proteins with preferences for high curvatures. We believe that a deeper understanding of curvature sensing requires the systematic characterisation of a larger pool of structurally different lipid-binding domains with different curvature preferences so that the influence of these factors can be studied separately. We believe our method can provide qualitative data on curvature preferences of a large number of lipid-binding domains and pave the way towards a better understanding of curvature sensing.

The reliance on nanomolar concentrations of protein for our method is, at the same time, an advantage and a caveat. While low protein concentrations can reduce the chance of membrane deformation by curvature sensors, it also reduces the chance to detect binding for domains with low affinity for membranes. For the lipid-binding domains used in this study, affinities for lipid membranes of 10nM-10μM have been reported (Frech et al., 1997; Uezu et al., 2011; Stahelin et al., 2002; Suetsugu et al., 2006; Dries and Newton, 2008; Ye et al., 2016; Sorre et al., 2012). The low nanomolar concentrations of protein used in this study are in a similar range thus allowing us to detect binding. It is however important to note that affinities of lipid-binding domains for membranes is extremely dependent on lipid composition, and in some cases curvature. Here, we used brain lipid extract as a membrane mimic that contains a variety of lipids. In cases where the lipid preference for specific membrane binding domains was available, the brain lipids have been supplemented with specific lipids to increase the affinity. Specifically optimising the lipid composition for each domain therefore allowed us to detect protein binding to liposomes despite the apparent discrepancy compared with published Kd.

As for any membrane-protein interaction study, different membrane compositions should be tested. Critically, as we show for Endophilin N-BAR, curvature preferences can be masked by strong attractive forces and therefore, a careful testing of lipid compositions might be required to correctly assign a curvature preference. An improved version of a NanoSight instrument we used in this study allows the use of a sample exchanger with automated sample injection that could streamline the screening of multiple lipid compositions and lipid binding domains.

In addition to curvature preference, we could also quantitatively analyse the vesiculation properties of the ENTH domain of Epsin. Crucially, the sensitivity of our method allowed us to discover that ANTH domains can also vesiculate membranes. We believe that vesiculation by ANTH domains has not been detected with the centrifugation-based assay previously used due to the differences in sizes of vesicles generated and the different vesiculation efficiency between ENTH and ANTH domains. We envisage that other lipid binding domains that can deform membranes via hydrophobic insertions may also display some vesiculation activity. The simplicity and resolution of our setup will allow researchers to probe this phenomenon in greater detail.

Using our method, we discovered that the PH domain of AKT and the FYVE domain of HRS preferentially bind to highly curved membranes. HRS is an endosomal protein and the preference for small liposomes fits with its cellular localisation and function (Kutateladze, 2006; Lawe et al., 2000). However, in the case of AKT, the explanation is less straightforward as AKT is found both on endosomes and on the plasma membrane (Andjelković et al., 1997; Schenck et al., 2008). It is tempting to suggest that the curvature preference of AKT may be part of a mechanism that allows this protein to differentially trigger signalling from membrane regions with high or low curvature. Further research will be necessary to understand the mechanism of this curvature preference and its biological implications.

Finally, our method allowed us to take a fresh look at the relationship between curvature sensing and curvature generation using Endophilin N-BAR as a model. The inverse correlation between membrane charge and curvature sensitivity further supports the idea that sensing and generation are two extremes of a continuum, *i*.*e*. the same protein domain can act both as a curvature sensor or a curvature generator depending on strength of its interaction with the membrane and its concentration. Furthermore, our results show that deletion of the Endophilin amphipathic helix H0 renders the protein primarily a curvature sensor. These results are in line with a recent study showing that the curvature preference of Endophilin in conditions of low protein concentration is primarily driven by low dissociation rates from smaller liposomes (Jin et al., 2022). Taken together, these results suggests that, at least in the case of Endophilin, the BAR domain is the primary sensor and the amphipathic helix is primarily a curvature generator. If this also applies to other lipid binding domains or if it is an idiosyncrasy of BAR domain proteins should be investigated further.

In conclusion, the method we present here is a flexible and generally applicable method that has the potential to significantly advance our understanding of the chemistry of membrane-protein interactions.

## Materials and Methods

### Materials

Lipids used are Brain Extract Folch fraction I (Sigma) (FolchS), Polar Brain Extract (Avanti) (FolchA), POPC (Sigma), brain PS (Sigma), brain PE (Sigma), cholesterol (Sigma), brain PI(4,5)P_2_ (Avanti), PI(3,4,5)P_3_-18:1 (Avanti), PI(3)P-16:0 (Sigma), PMA (Sigma), and **DiOC**_**18**_**(3)** (Thermo Fisher Scientific).

NS buffer contains 20mM HEPES pH7.4, 100mM NaCl, 0.5mM TCEP. For protein purification, following buffers were used: IMAC-L (20mM Tris pH8.0, 200mM NaCl, 50mM Imidazole, 0.5mM TCEP), IMAC-E (20mM Tris pH8.0, 200mM NaCl, 250mM Imidazole, 0.5mM TCEP), IEX-A (20mM Tris pH8.0, 0.5mM TCEP), IEX-B (20mM Tris pH8.0, 500mM NaCl, 0.5mM TCEP), GEF (20mM HEPES pH7.4, 150mM NaCl, 0.5mM TCEP).

### Liposome preparation

Lipid stocks in chloroform were mixed in a glass vial. The solvent was evaporated against the walls of the vial using an argon stream. The dried lipid film was then placed for 30min in a desiccator to completely evaporate remaining organic solvents and water. For long-term storage, the vial was filled with argon gas and stored at -20°C. Lipids were resuspended at a concentration of 0.25mg/ml in NS buffer by rolling for 1-2h at room temperature. The solution was vortexed twice for 20s each during this time. Liposomes were extruded using 800nm, 200nm, 100nm and 50nm Whatman Nucleopore Polycarbonate filters in an Avanti Mini Extruder. Fresh liposomes were kept at room temperature and used within 24h.

For sizing a fluorescent liposome subpopulation, liposomes were made of FolchS (non-fluorescent), and 1% or 10% **DiOC**_**18**_**(3)** was added to the large and small fluorescent liposomes respectively. For measurements of the correction factor, FolchS with 10% **DiOC**_**18**_**(3)** was used. Liposomes used for measurements of curvature sensitivity of Amphiphysin contained 38% POPC, 25% PE, 20% PS, 2% PI(4,5)P_2_ and 15% Cholesterol (values given in molar percentages). A 1:1 mix of Sigma and Avanti brain extract lipid (FolchSA) was used for FCHo2, IRSp53 and Endophilin ΔH0 experiments. For other lipid binding domains, FolchS was spiked with 2% PIP3 (AKT PH), 2% PI(3)P (HRS FYVE), 1% PMA (PKCß2 C1B) or 2% PI(4,5)P_2_ (Talin1 FERM). For vesiculation with ENTH and ANTH domains, liposomes were made of FolchS with 2% PI(4,5)P_2_ and 1% **DiOC**_**18**_**(3)**.

### Cloning

Following constructs were used: rat Endophilin A2 N-BAR (amino acids 1-247), rat Endophilin A2 BAR ΔH0 (25-247), mouse FCHo2 BARX (1-327), human IRSp53 BAR (1-250), human Amphiphysin N-BAR (1-252), human Amphiphysin ΔH0 (25-252), human Talin1 FERM (1-401), human AKT PH (1-164), human PKCb2 C1B (91-161), human HRS FYVE (149-230), human Epsin ENTH (1-164), human PiCALM ANTH (87-289), human Hip1R ANTH (1-161).

Constructs were cloned using Fragment Exchange (FX) cloning(Geertsma and Dutzler, 2011). Monomeric enhanced superfolder GFP (mesfGFP)(Zhang et al., 1996; Zacharias et al., 2002; von Stetten et al., 2012b) fusion constructs used in curvature-sensing experiments were cloned with an N-terminal His10-mesfGFP-linker-3C or a C-terminal 3C-mesfGFP-His10 tag (3C = PreScission cleavage site). For measurements of vesiculation, ENTH and ANTH domains were expressed with an N-terminal His10-SUMO tag.

### Recombinant protein expression in E. coli

Vectors containing the gene of interest under the control of the T7 promoter were transformed in BL21(DE3) cells (Thermo Fisher Scientific) and plated on TYE-agar containing the corresponding antibiotic for selection. The next day, colonies were inoculated in 50ml 2xTY. After a few hours, this 20ml preculture was added to 1l 2xTY and cells were grown until OD_600_ reached 0.8-1. Protein expression was then induced by addition of 160μM IPTG overnight at 18°C. For small-scale protein expression, the protocol was similar except that 1ml preculture was added to 50ml 2xTY.

### Small-scale protein purification

50ml cultures were harvested by centrifugation for 15min at 3000g. Pellets were resuspended in 3ml IMAC-L containing lysozyme and EDTA-free Proteoloc Protease Inhibitor cocktail (Expedeon) and incubated for 10min at 4°C. Cells were lysed by sonication using a Microson Ultrasonic cell disruptor with a micro tip (Misonix incorporate). Unbroken cells and debris were pelleted for 5min at 20000g. The supernatant was transferred in a fresh tube. After addition of DNaseI 1mM MgCl_2_ and 200μl 50% TALON^R^ beads slurry (Clontech), the cell lysate was incubated at 10min at 4°C on a rolling shaker. Beads were washed with 10ml IMAC-L, 1ml IEX-B (high-salt wash), 15ml IMAC-L. Protein was eluted with 1ml IMAC-E and further purified by size exclusion chromatography as described below.

### Large-scale protein purification

Large-scale protein purification was generally realised in three steps, an affinity capture using His-tag, followed by an ion exchange column and finally size exclusion chromatography. Cultures were harvested by centrifugation for 15min at 4200g. Pellets were resuspended in IMAC-L containing lysozyme and EDTA-free Proteoloc Protease Inhibitor cocktail (Expedeon) and incubated for 10min at 4°C. Cells were lysed by sonication using a Sonics VC 750. After addition of DNaseI and 1mM MgCl2, unbroken cells and debris were pelleted 15min at 40000g. The supernatant was loaded onto HisTrap™HP column (GE Healthcare). HisTrap column were washed with IMAC-L and IEX-B, then protein was eluted with IMAC-E.

Depending on the pI of the protein, anion exchange (HiTrap™Q, GE Healthcare) or cation exchange (HiTrap™SP, GE Healthcare) chromatography was used. Prior to loading on ion exchange column, NaCl and imidazole were diluted out in IEX-A. A NaCl gradient ranging from 100mM to 500mM NaCl was run on ÅKTA Purifier 10 system.

Protein domains expressed with a SUMO tag were cleaved by SENP1 protease.

For size exclusion chromatography, either a Superdex™75 or a Superdex™200 column (GE Healthcare) was used depending on the size of the protein. The amount of protein determined which size of column was used: 10/30, HiLoad 16/60 or HiLoad 26/60. Size exclusion chromatography was run in GEF buffer. Protein was concentrated using Amicon Ultra centrifugal filter units (Merck Millipore). Protein was then aliquoted, flash-frozen in liquid N_2_ and stored at -80°C.

### Vesiculation

For vesiculation experiments, liposomes at 0.25mg/ml lipids were incubated for 30 minutes at 37°C in a PCR machine with the indicated concentration of non-fluorescent protein. For EM, the sample was used pure. For measurements, the sample was diluted 500 times.

### Electron microscopy

For EM, formvar/carbon, glow-discharged grids were immediately placed on a drop containing liposome samples and incubated for 2 minutes. Grids were then dried on a filter paper, stained twice for 20s each with 3% uranyl acetate, rinsed and dried overnight. For measurements with Endophilin N-BAR and BAR ΔH0, 2μM protein and 1.25μg/ml liposomes were used to keep the ratio of protein to lipid constant compared to samples used for single particle tracking

The fraction of large liposomes (diameter >100nm) showing protrusions in each electron micrograph was quantified. ANOVA with Bonferroni correction for multiple comparisons was used to compare means and calculate p-values.

### Instrumentation and measurements

Measurements were done on a NanoSight LM10 (Malvern) equipped with a sCMOS camera and a Harvard Apparatus syringe pump. A 488nm laser together with a 500nm long pass filter was used for green fluorescence. To check the calibration, 100nm and 216nm NIST (National Institute of Standards and Technology) traceable calibration beads (3000 Series Nanosphere^R^ Size Standards (Thermo Fisher Scientific)) were diluted in water. Movies were recorded without pump flow and particles were tracked using the company’s software. Results used were FTLA (Finite Track Length Adjustment) corrected using the proprietary algorithm. For comparison with Dynamic Light Scattering, a W130i DLS system (AvidNano) was used.

For measurements of curvature sensitivity, liposome solutions were diluted to reach final 2-8 x 10^8^ particles/ml as recommended by the manufacturer. This corresponded to final lipid concentrations of about 1.25μg/ml for unextruded or 800nm extruded liposomes and 0.125-0.25μg/ml for 200nm or 50nm extruded liposomes. Fluorescent protein concentration was 1-3nM. Liposomes were first diluted in NS buffer, then protein was added. After mixing, the sample was loaded onto the NanoSight. Recordings were made under flow from the syringe pump (setting 50) to reduce photobleaching. 120s long movies were recorded at 25fps using appropriate camera settings to maximise signal/background ratio. Particles were tracked using the company’s software and their size calculated based on their Brownian motion (ISO 19430:2016). Raw, non-FTLA corrected data was used due to sample heterogeneity.

### Data Analysis

Single particle tracking and data processing was carried out by the NanoSight NTA software version 3.1 (Malvern). Raw size distributions binned in 5nm bins were extracted directly from the Nanosight NTA software. For diffraction measurements of 50nm extruded liposomes, a correction factor (CF) was subsequently applied to account for the discrepancy between recordings using diffraction or fluorescence due to the weakly diffracting liposomes. The CF curve was calculated by measuring the difference between the distribution curves of fluorescent 50nm liposomes in diffraction (Diff) and fluorescence (Fluo) modes. The CF for each bin was obtained by applying the formula CF = (Fluo – Diff)/Diff (Supplementary fig. S1d-f).

Smoothing of raw curves was performed in Excel using a 7 datapoint running window. Box plots were generated by using the mode for the middle line and 50% of the data on each side of the mode were used for boundaries. A typical run measures between 3000 and 8000 particles. For such large n, even small differences between replicates, that are not biologically relevant, become statistically significant. Therefore, for statistical differences, mode values for each experimental replicate (i.e. the curvature preference for a specific protein) were used as single data points. ANOVA with Bonferroni correction for multiple comparisons were used to calculate p-values (significance levels: ^*^p < 0.05, ^**^p < 0.01, ^***^p < 0.001, ^****^p < 0.0001). Statistical analyses were performed in Graphpad prism 7.0.

### Data availability

The authors declare that all data supporting the findings of this study are available within the paper and its supplementary information files.

## Author contributions

A.C. performed all experiments. A.C. and L.A.-S. designed research and collected EM images. L.A.-S. performed data processing and statistical analysis. L.A.-S. and H.T.M. supervised the project. L.A.-S., A.C. and H.T.M. wrote the manuscript.

## Acknowledgments

We would like to thank our colleagues at the MRC LMB scientific facilities, including Chris Johnson and Stephen McLaughlin for biophysics; Shaoxia Chen and team for EM; Mark Skehel and team for Mass Spectrometry. We are grateful for Vladan Martinović (MRC-LMB, Cambridge, UK) for the purified Hip1R and PiCALM ANTH domains. We thank Rohit Mittal and David Paul for critical reading of this manuscript. This work was supported by the Medical Research Council (grant number MC_U105178795). LA-S is supported by a HiLIFE start-up grant and the Academy of Finland (Research Fellow). L.A.-S. was an EMBO Long Term fellow supported by Marie Curie Actions.

