## Supplemental Figures for "A single particle analysis method for detecting membrane remodelling and curvature sensing"

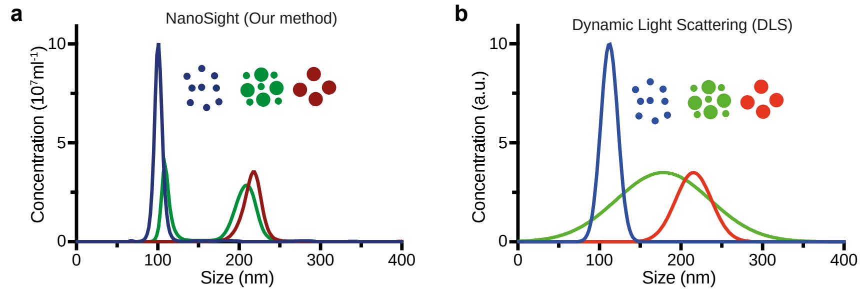


c

| beads | NanoSight | DLS |
|---|---|---|
| 100nm | 100±5nm | 110±10nm |
| 216nm | 217±11nm | 214±22nm |

| beads | NanoSight | DLS |
|---|---|---|
| Mix of 100, 216nm | 2 peaks 109±5nm 208±13nm | 177±58nm |

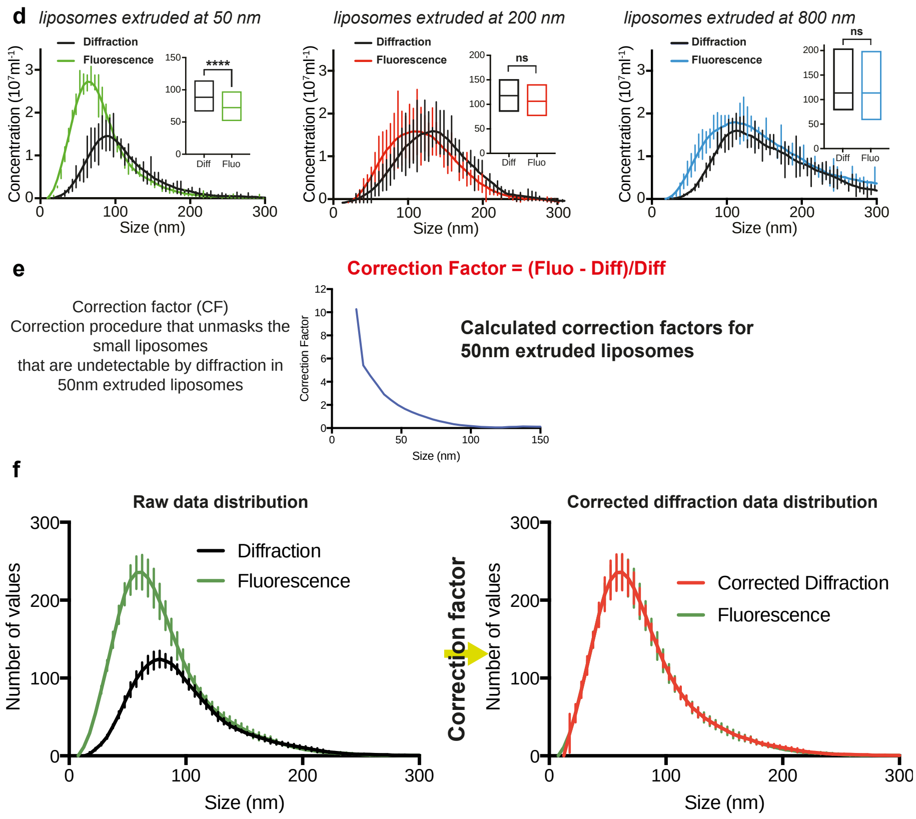


**Figure S1 – Data processing, visualisation and correction**

**a-c**. Comparison of sizing of 100 nm (blue), 216 nm (red) beads or a mix (green) by NanoSight (a) or DLS (b) and corresponding bead size measurements (c). Results are indicated as mean ± standard deviation of bead diameters.

**d**. Comparison of size distributions of fluorescent liposomes using diffraction or fluorescence measurements shows that smaller liposomes are better detected by fluorescence than by diffraction. This is especially obvious with liposomes extruded at 50 nm. Boxplots depict mode values (middle line) -/+ 50% data on each side of the mode (indicated by bottom and top lines). n =3, ns = non-significant, p<0.0001 (****). Two tailed Student's t-test.

**e-f**. To correct for the non-detectable liposomes from measurements using diffraction, a correction factor is applied. Application of the correction factor on the diffraction data (f, right, black curve) results in a corrected diffraction size distribution (f, left, red), which overlaps well with the size distribution obtained by fluorescence (f left, green).

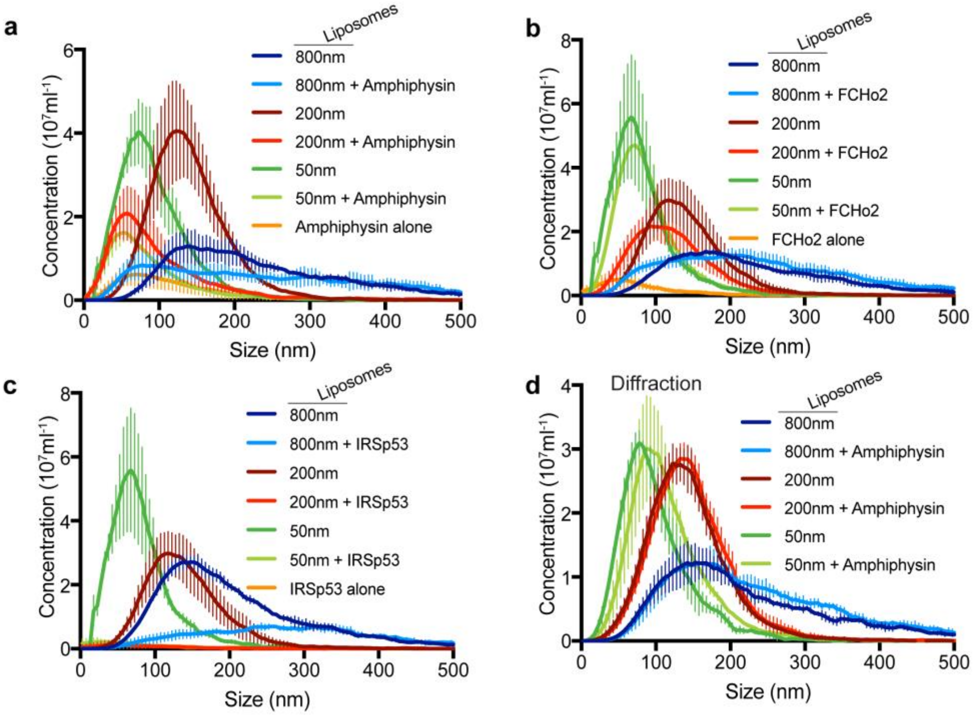


**Figure S2 - Raw size distributions for measurements using BAR domains**

**a-c**. Size distributions of liposomes extruded at 800 nm (dark blue), 200 nm (dark red) or 50 nm (dark green) with corresponding size distributions of these liposomes detected by the presence of bound protein (light blue, light red, light green) using Amphiphysin N-BAR (a), FCHo2 F-BAR (b) or IRSp53 I-BAR (c). Signal from protein in the absence of liposomes is shown in orange.

**d**. Size distributions of the total liposome population (detected by diffraction) in the absence (darker colours) or presence (lighter colours) of Amphiphysin N-BAR shows that no large-scale remodelling of liposomes occurs.

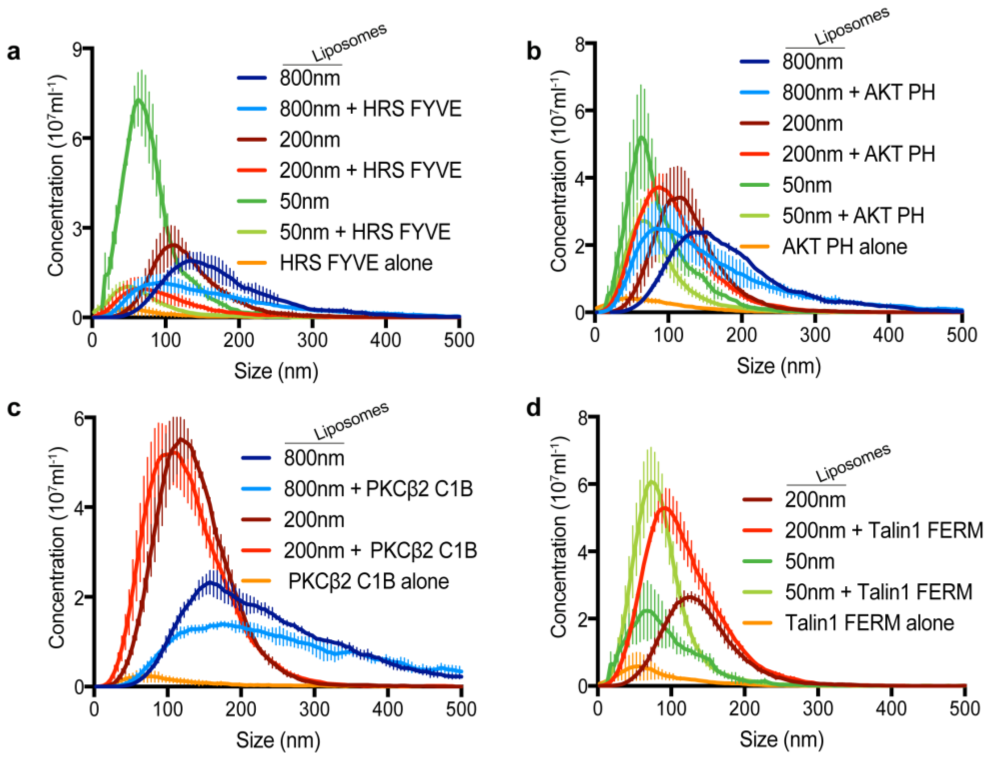


**Figure S3 - Raw size distributions measurements of diverse lipid-binding domains**

**a**. Size distributions of liposomes extruded at 800 nm (dark blue), 200 nm (dark red) or 50 nm (dark green) with corresponding size distributions of these liposomes detected by the presence of bound HRS FYVE (light blue, light red, light green). Signal from HRS FYVE in the absence of liposomes is shown in orange.

**b**. Size distributions of liposomes extruded at 800 nm (dark blue), 200 nm (dark red) or 50 nm (dark green) with corresponding size distributions of these liposomes detected by the presence of bound AKT PH (light blue, light red, light green). Signal from AKT PH in the absence of liposomes is shown in orange.

**c**. Size distributions of liposomes extruded at 800 nm (dark blue) or 200 nm (dark red) with corresponding size distributions of these liposomes detected by the presence of bound PKCβ2 C1B (light blue, light red). Signal from PKCβ2 C1B in the absence of liposomes is shown in orange.

**d**. Size distributions of liposomes extruded at 200 nm (dark red) or 50 nm (dark green) with corresponding size distributions of these liposomes detected by the presence of bound Talin1 FERM (light red, light green). Signal from Talin1 FERM in the absence of liposomes is shown in orange.

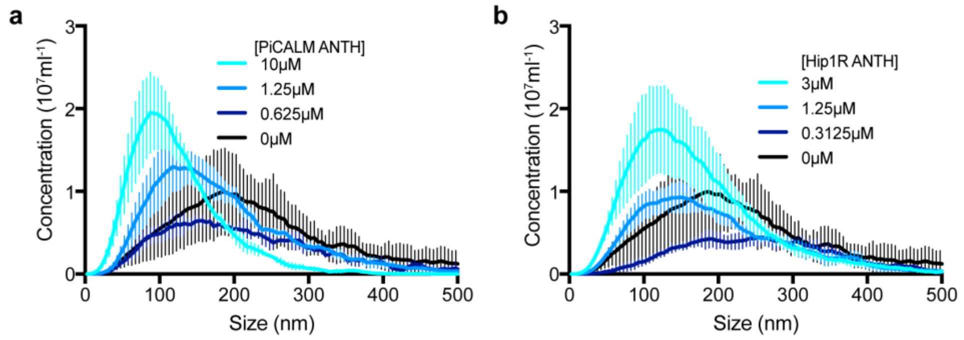


**Figure S4 – Detecting membrane vesiculation**

**a, b**. Size distributions of liposomes showing a dose dependent vesiculation by PiCALM ANTH (a) and Hip1R ANTH (b).

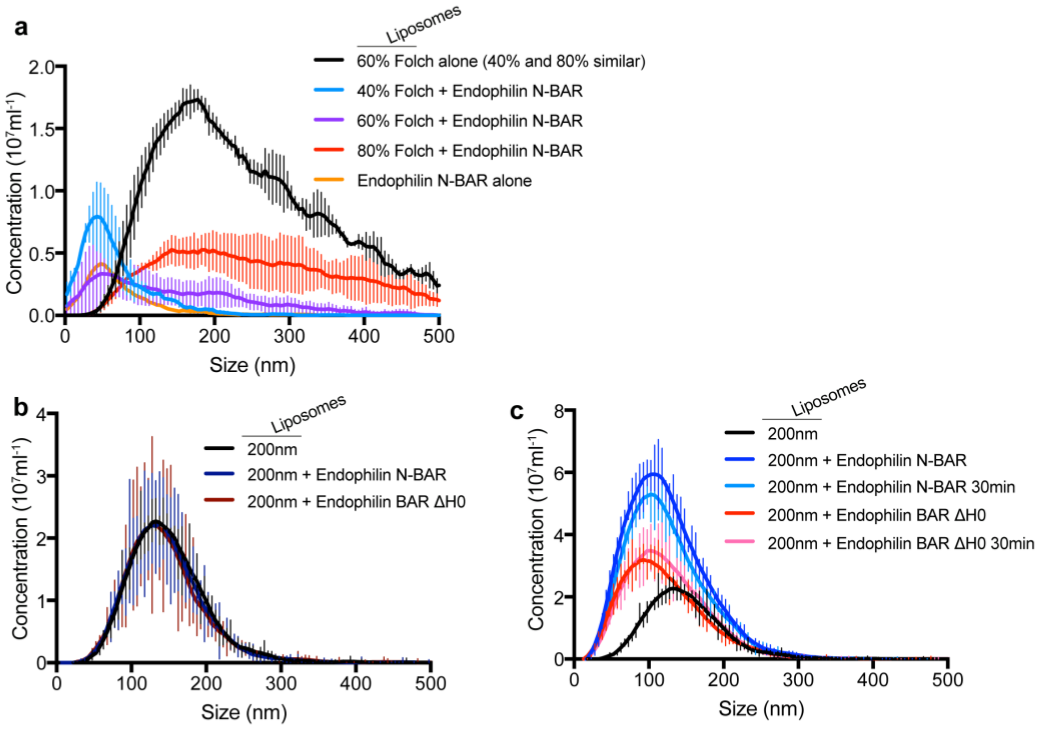


**Figure S5 - Mechanism of curvature sensing by Endophilin**

**a**. Raw size distributions for measurements of Endophilin N-BAR-GFP on liposomes with increasing concentrations of Folch lipids.

**b**. Similar size distributions of liposomes before (black) and after addition of Endophilin N-BAR (blue) or Endophilin BAR ΔH0 (red) shows that no vesiculation is taking place.

**c**. Longer incubation (30 minutes) with Endophilin N-BAR (light blue) or BAR ΔH0 (pink) does not change the size of the protein-bound liposomes compared to immediately after mixing protein and liposomes (dark blue for Endophilin N-BAR, red for Endophilin BAR ΔH0) indicates that neither of the protein vesiculates liposomes.

**Table S1 – Results from the screening of lipid binding domains**

| Domain | Gene | Lipid composition | Binding | Curvature sensing |
|---|---|---|---|---|
| **PH** | AKT1[33] | FolchS + 2% $PIP_3$ | + | High curvatures |
| **PTB** | DAB2[34,35] | FolchS + 2% $PI(4,5)P_2$ | + | Not tested |
| **GRAM** | OXR1 | FolchSA | - | Not applicable |
| **C1**[36] | PRKCB2 | FolchS + 1% PMA | + | Curvature insensitive |
| **C2** | PLA2G4A[37] | POPC (+ $CaCl_2$) | - | Not applicable |
| **C2** | SYT1[38,39] | FolchSA + 10% $PI(4,5)P_2$ or $PIP_3$ (+ $CaCl_2$) | - | Not applicable |
| **FYVE**[28,40] | HGS | FolchS + 2% PI(3)P | + | High curvatures |
| **FERM**[41] | TLN1 | FolchS + 2% $PI(4,5)P_2$ | + | Curvature insensitive |
