## Supplementary material for "A single particle analysis method for detecting membrane remodelling and curvature sensing": Suplemental Note

### Supplementary Note 1

### - Introduction to the Nanosight technology -

The instrument, NanoSight LM10 (Malvern), used in this study is based on a specially designed viewing unit mounted on a conventional upright microscope with a long working distance 20x objective and equipped with a high sensitivity CMOS camera (Fig. 1a). The viewing unit consists of an imaging chamber (12mm Ø) mounted on a laser block. The unit has ports for sample in/output and for the temperature sensor (Fig. 1b).

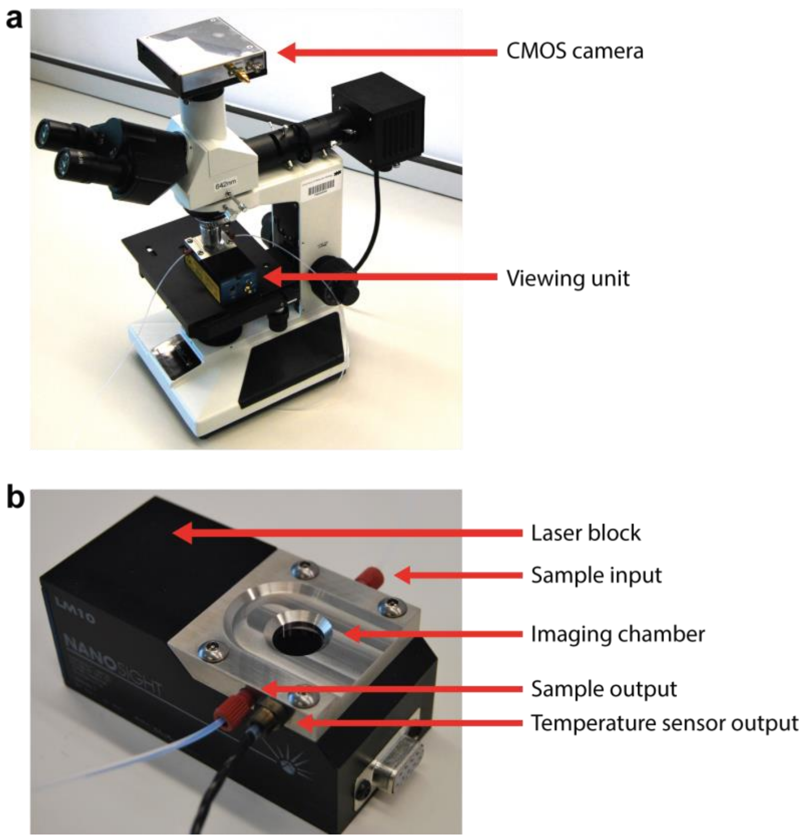


*Figure 1 – Overview of the NanoSight.* ***a****. The Nanosight is based on a conventional upright microscope.* ***b****. Close up picture of the Nanosight viewing unit.*

The sample is injected into the viewing unit using a syringe pump (Fig. 2a) and imaged in a specially designed glass chamber where the bottom surface, called the optical flat, is coated with a metallised surface to reduce background (Fig. 2b). Light from the integrated laser passes through the non-coated window in the optical flat and gets refracted upon reaching the liquid sample, forming a beam through the sample (Fig. 2c). Figure 2d shows a low magnification view of the flow chamber using a 4x objective. The flare on the right represents the point at which the laser exits the optical flat, the vertical line to its left is the boundary between the coated and uncoated glass surface. Measurements are taken in an 80 µm x 100 µm x 10 µm observation volume (indicated by the red box in Fig. 2d) next to the boundary line on the coated surface, where the laser is brightest. As the laser beam goes through the sample, it quickly loses brightness.

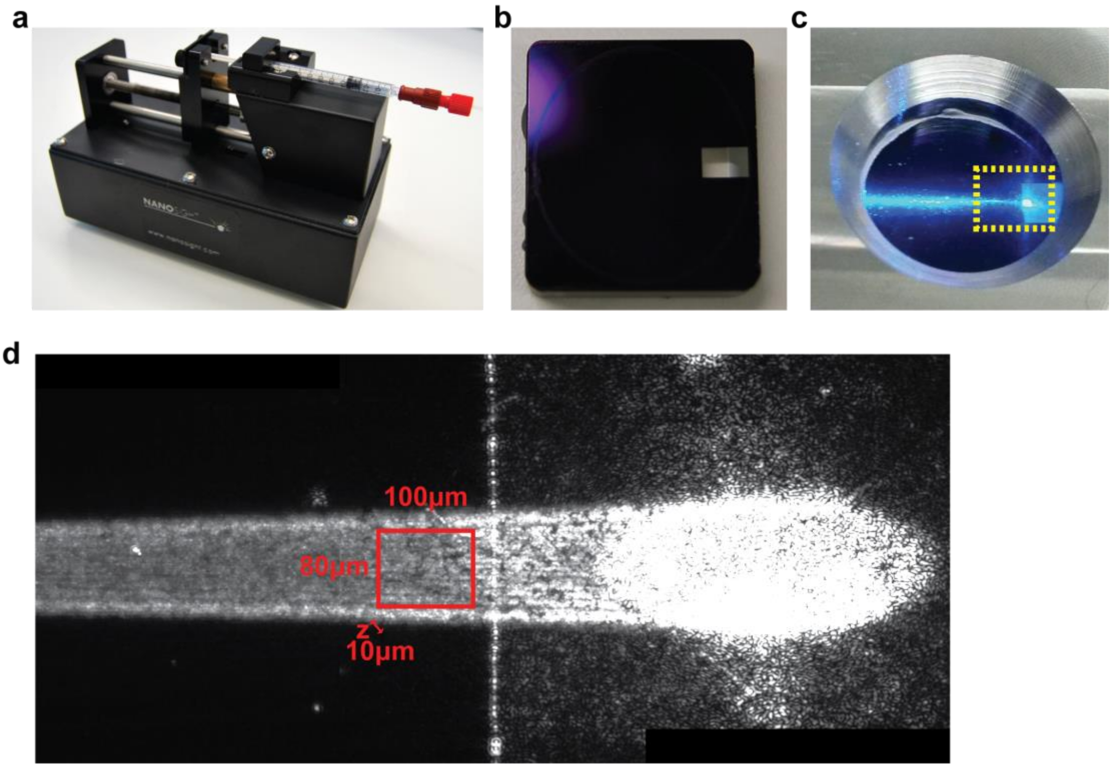


*Figure 2 – **a**. The syringe pump used to inject samples into viewing unit. **b**. the bottom part of the chamber (optical flat) has a special coating to reduce background. Note the opening on the right where the laser enters the chamber. **c**. View of the laser beam passing through the imaging chamber. **d.** Low magnification view of the area marked on yellow in (c). The red square marks the approximate region where videos for tracking are captured.*

Measurements consist of 120 s movies recorded using the high-sensitivity CMOS camera operating at 25 frames per second. Particles are detected using diffraction of the incident laser (Fig. 3a,c), whereas protein-bound liposomes are identified as fluorescent particles using a long pass emission filter (Fig. 3b,d). The lack of background fluorescence derived from liposomes is illustrated by comparing images of a sample of regular liposomes detected using diffraction (Fig. 3f) and under fluorescence (Fig. 3g). In addition, signal from protein alone (here FchO2 as an example) is also shown (Fig. 3h).

After background subtraction, the centre of mass of each particle is determined. The threshold for detection of particles can be adjusted depending on the intensity of the particles present in the sample. Particles are then automatically tracked and their size calculated based on their Brownian motion. The average displacement of all particles across the field of view is measured and used to subtract the flow generated by the pump in the Brownian motion calculation. The algorithm used for particle tracking is described in ISO 19430:2016. Figure 3i shows an example of temporally colour-coded particles moving along the filed of view.

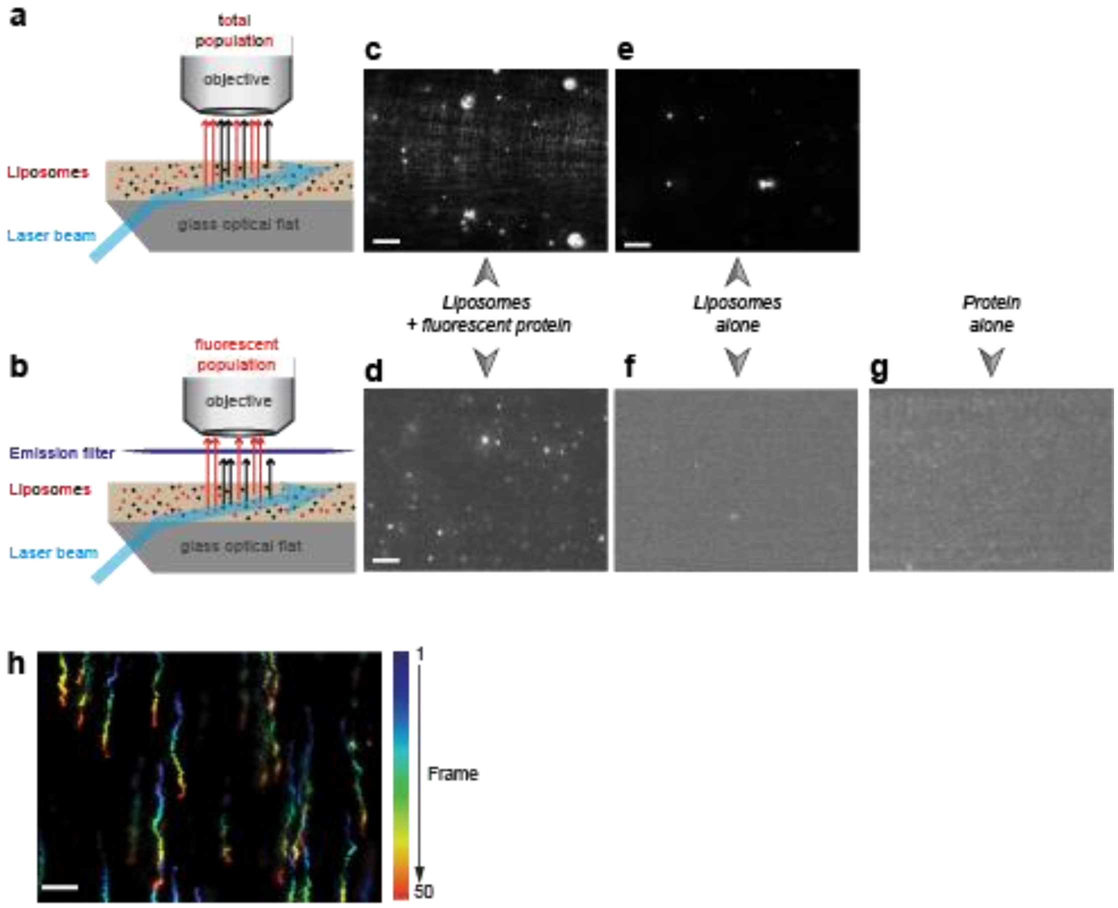


*Figure 3 – Particle detection and tracking.* ***a,b****. Particle passing through the imaging chamber and hit by the incident laser beam can be detected by diffraction (a) or fluorescence emission (b).* ***c,d****. Typical images of liposomes and fluorescent proteins captured diffraction(c) and fluorescence (d).* ***e,f.*** *Typical images of liposomes alone captured under diffraction (e) and fluorescence (f).* ***g.*** *Typical image of protein alone (here FChO2) imaged by fluorescence.* ***h****. Temporally colour-coded image of particles crossing the Nanosight field of view. Time intensity projection of 50 frames (2 s). Each frame of the video was assigned a colour as shown in the legend (right) before generating the projection image. Scale bar = 10µm*

Although fluorescence can be used to detect particles, quantifying its intensity to calculate the fluorescent-protein coverage on each liposome is limited by intrinsic characteristics of the machine. The most important limiting factor for fluorescence measurement is that the illumination is not even across the

Nanosight field of view. As can be seen in Fig. 2d, illumination varies both in "x", along the axis of the laser beam, due to scattering of the light as the laser crosses the sample as well as in "y" due to imperfect glass surface where the laser is diffracted. Moreover, as particles move freely in three dimensions and are imaged in widefield mode, movements in z will also result in varying detected intensities as particles out of focus will still appear on the image and have a clear centre of mass to be able to be tracked. In addition, fluorescent signals decay as the particles flow along the chamber due to bleaching.
